# Comprehensive Infectome Analysis Reveals Diverse Infectious Agents with Zoonotic Potential in Wildlife

**DOI:** 10.1101/2025.01.09.632079

**Authors:** Xin Hou, Jing Wang, Da-Xi Wang, Ye Chen, Xin-Xin Li, Wei-Chen Wu, Shi-Jia Le, Shi-Qiang Mei, Han-Lin Liu, Yun Feng, Li-Fen Yang, Pei-Bo Shi, Zi-Rui Ren, Lin Yang, Hai-Jian He, Wan-Jie Song, Jie Chen, Meng-Meng Zhu, Zhi-Wen Jiang, Xing-Mei Li, Mei-Rong Li, Deyin Guo, Edward C. Holmes, Zi-Qing Deng, Mang Shi

## Abstract

Understanding wildlife-pathogen interactions is crucial for mitigating zoonotic risk. Through meta-transcriptomic sequencing we profiled the infectomes of 1,922 samples from 67 mammalian species across China, uncovering a remarkable diversity of viral, bacterial, fungal, and parasitic pathogens. Of the 195 pathogens identified, 62 were novel, including a bi-segmented coronavirus in diseased lesser pandas, which we propose represents a new genus – Zetacoronavirus. The orders Carnivora and Rodentia exhibited the highest pathogen diversity and were implicated in numerous host-jumping events. Comparative analysis of diseased versus healthy animals revealed a trend of higher pathogen loads in the former, with possible differences in tissue tropisms. In total, 48 zoonotic and 17 epizootic pathogens were identified, with frequent cross-species transmission, emphasizing the potential for emerging public health threats. This study highlights the urgent need for wildlife pathogen surveillance to inform proactive disease management strategies.

## Introduction

Mammals host a diverse array of viruses, bacteria, eukaryotic parasites and fungi that can cause disease in humans and other animals^1–7^. Cross-species transmission enables these pathogens to occasionally jump to humans, sometimes resulting in localized outbreaks or even large-scale epidemics and pandemics. To trace the origins of particular viral pathogens, and to help prevent future emergence events, considerable attention has been devoted to discovering viruses in wildlife using metagenomic approaches^8–14^. While most of the viruses discovered primarily affect a single wildlife host species, others have crossed species boundaries, or even emerged in different mammalian orders, suggesting that they might be host generalists with the potential to emerge in multiple hosts, perhaps including humans^15,16^.

Although metagenomics has brought a new perspective on the diversity of the virosphere, the understandable emphasis on viral infections has meant that less is known about the roles played by bacteria and eukaryotic pathogens in disease emergence. However, reports of bacterial and fungal infections associated with human mortality, including those carrying antimicrobial resistance genes, highlight the need for a broader perspective on disease emergence^17^. Meta- transcriptomic (i.e., total RNA) sequencing, which can simultaneously reveal viruses, bacteria, and eukaryotic pathogens, provides an efficient and increasingly cost-effective way to obtain a comprehensive understanding of the total “infectomes” of wildlife species^18,19^. Herein, we conducted an extensive individual-level infectome analysis in a diverse array of mammals from China, including common and rare species, domestic and wild animals, as well as diseased and healthy individuals. For many species, including primates (e.g., macaques and ring-tailed lemurs), artiodactyls (e.g., sika deer and muntjacs), and perissodactyls (e.g., horses and donkeys), such analyses are rare. Our particular aim was to systematically characterize the pathogen composition in a wide range of mammalian species, explore their interactions and cross-species transmission, and provide crucial insights for the early warning and control of human infectious diseases.

## Results

### The mammalian total infectome

Between 2018 to 2024 we collected 1,922 samples from 67 species across 11 mammalian orders in 20 Chinese provinces, comprising 1,283 diseased and 639 healthy samples (Fig. 1a, Extend Data Fig 1 and Supplementary Table 1). This sampling included rare animal species such as tufted capuchins (*Sapajus apella*), snub-nosed monkeys (*Rhinopithecus roxellana*) Chinese water deer (*Hydropotes inermis*), six-banded armadillos (*Euphractus sexcinctus*), and caracals (*Caracal caracal*), which have not previously systematically examined for potential pathogens. Various organs relevant to disease outbreak, including liver, spleen, lung, kidney, intestine, and lymph nodes from diseased animals were collected. While faecal and respiratory swabs were obtained from healthy animals for comparison. Subsequently, we employed the meta- transcriptomic sequencing to profile the total infectome which led to the characterization of a whole spectrum of virus, bacteria, fungi, and parasitic pathogens within these samples. These include: (i) established pathogens, (ii) opportunistic pathogens, and (iii) previously undescribed microorganisms with pathogenic potential in mammals. Phylogenetic assessments and abundance estimations (RPM > 10) confirmed the presence of 105 RNA viral species, 42 DNA viral species, 34 bacterial pathogen species, ten fungal pathogen species, and four parasitic species (Fig. 1b, Extend Data Fig 2 and Supplementary Table 2). Among them, 62 novel pathogen species were discovered, comprising 58 species of viruses and four divergent members of the fungal genus *Pneumocystis*.

**Fig. 1.**
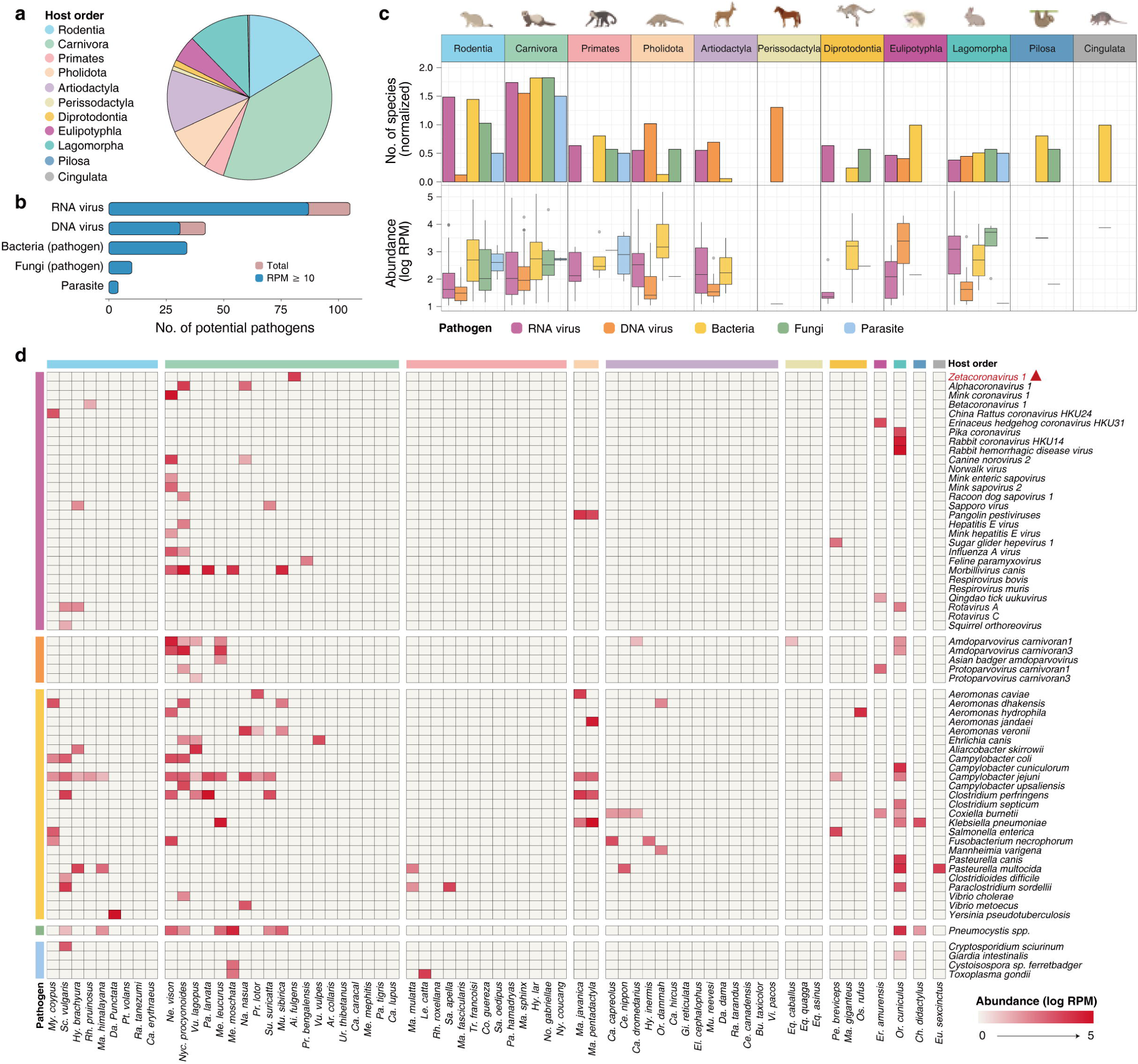
Total infectome characterization of mammalian wildlife. a, Number of samples collected at the level of mammalian order. **b,** Number of potential viral, bacterial, fungal and parasitic pathogens identified in this study. The abundance levels of 29 pathogens fell below our classification threshold (i.e., 10 RPM) for positivity and were excluded from further analysis. **c,** Pathogen richness and abundance at the animal order level. Pathogen richness was normalized based on sample size for comparison. **d,** Distribution and abundance of representative pathogens with potential link to disease. The relative abundance of pathogens in each library was calculated and normalized by RPM. To remove potential contamination, only pathogens with an abundance RPM ≥10 are shown. Pathogen groups and mammalian orders are shown by the colors on the heatmap.

**Fig. 2.**
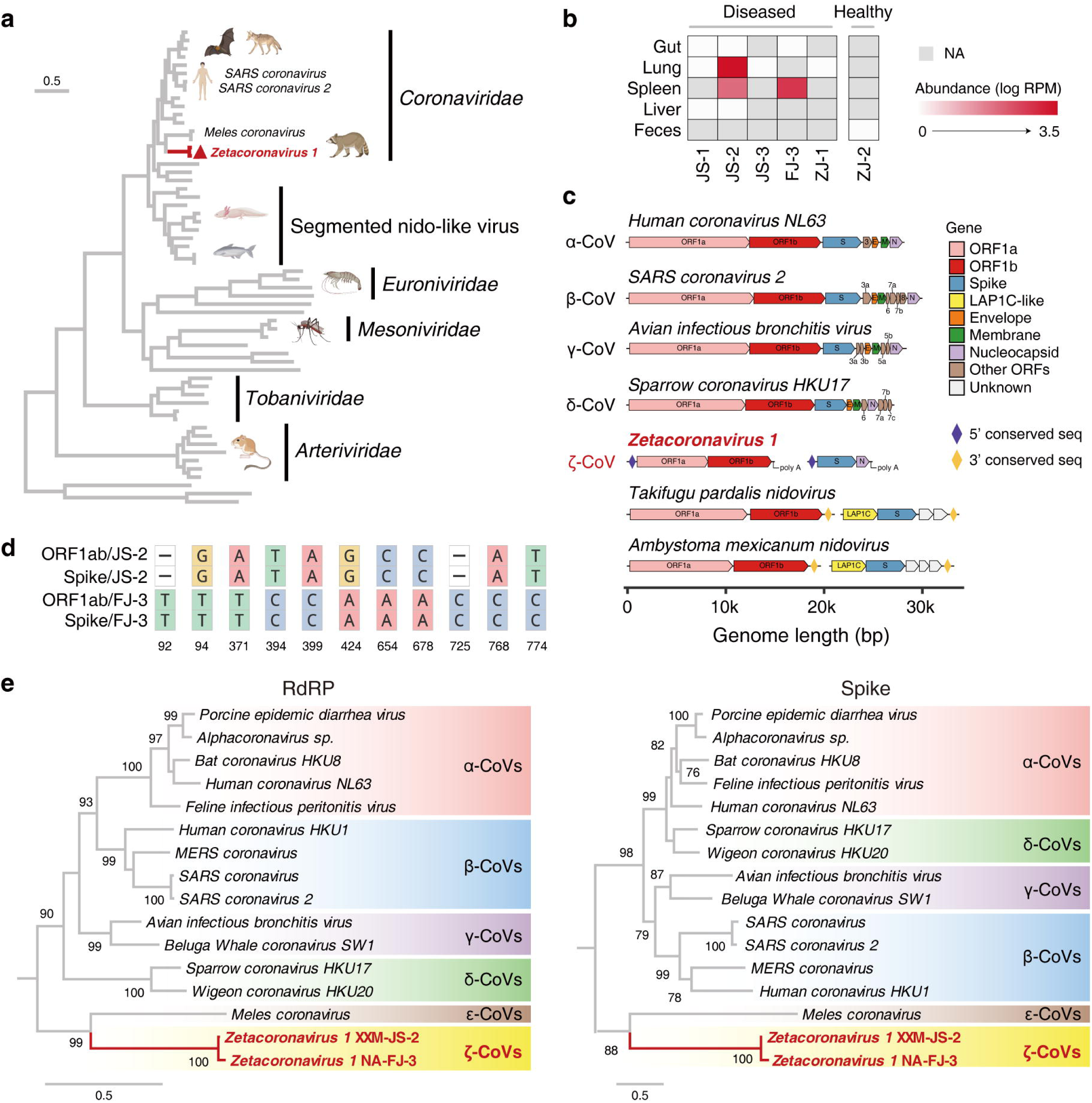
Novel pathogenic and segmented coronavirus in diseased lesser pandas. a, The phylogenetic position of the novel coronavirus – tentative species Zetacoronavirus 1 *–* identified in this study is depicted in the phylogenetic tree of the *Nidovirales* and marked by red bold font. **b,** The prevalence and abundance of Zetacoronavirus 1 in lesser pandas. “NA” denotes the sample types are not available for sample collection in individuals. **c,** The genomic structures of common coronaviruses, as well as Zetacoronavirus 1 and segmented Nido-like viruses. Zetacoronavirus 1 contains a conserved sequences of around 700bp in 5’ end of each segment, while the segmented Nido-like viruses contain an identical 129-nt trailing sequence in 3’ end of each segment. **d,** Single nucleotide polymorphisms (SNPs) of the conserved 700 bp sequences in the two strains of Zetacoronavirus 1. **e,** Phylogenetic tree of *Coronaviridae* using a maximum likelihood method based on RdRP and spike protein sequences. All trees were midpoint-rooted for clarity only, and the scale bar indicates 0.5 amino acid substitutions per site. Zetacoronavirus 1 is marked in red bold font.

Our standardized analysis, adjusted for variation in sample sizes, revealed that the mammalian order Carnivora harboured the most diverse and abundant pathogens, followed closely by the Rodentia (Fig. 1c, Extend Data Fig 2 and 3). While high viral diversity and abundance in Rodentia can likely be attributed to their often high population densities, the elevated viral diversity and abundance in Carnivora more likely reflects their unusually high population density in fur farms as opposed to natural environments. Among the pathogens identified, we found zoonotic viruses such as avian influenza viruses H5N1 and H5N6, *Alphacoronavirus 1*, *Norwalk virus*, *Rotavirus A*, *Hepatitis E virus* (HEV), and several viruses known to cause acute or severe diseases to mammals including *Morbillivirus canis*, *Rabbit hemorrhagic disease virus (RHDV)*, *Pangolin pestiviruses*, *Feline paramyxovirus, Respirovirus bovis and Respirovirus muris, Sapporo virus, Amdoparvovirus carnivoran1* and *Amdoparvovirus carnivoran3* (Fig. 1d, Extend Data Fig 4 and 5a). In addition to the well-known mammalian-infecting viruses mentioned above, several previously uncharacterized viral agents at high abundance (≥10 RPM) were identified in diseased animals and warrant further attention. Also of note was the identification of two novel sapoviruses in mink and raccoon dogs, as well as a novel strain of *Hepatitis E virus* – specifically *Mink hepatitis E virus* – in mink populations from Shandong and Hebei provinces.

**Fig. 3.**
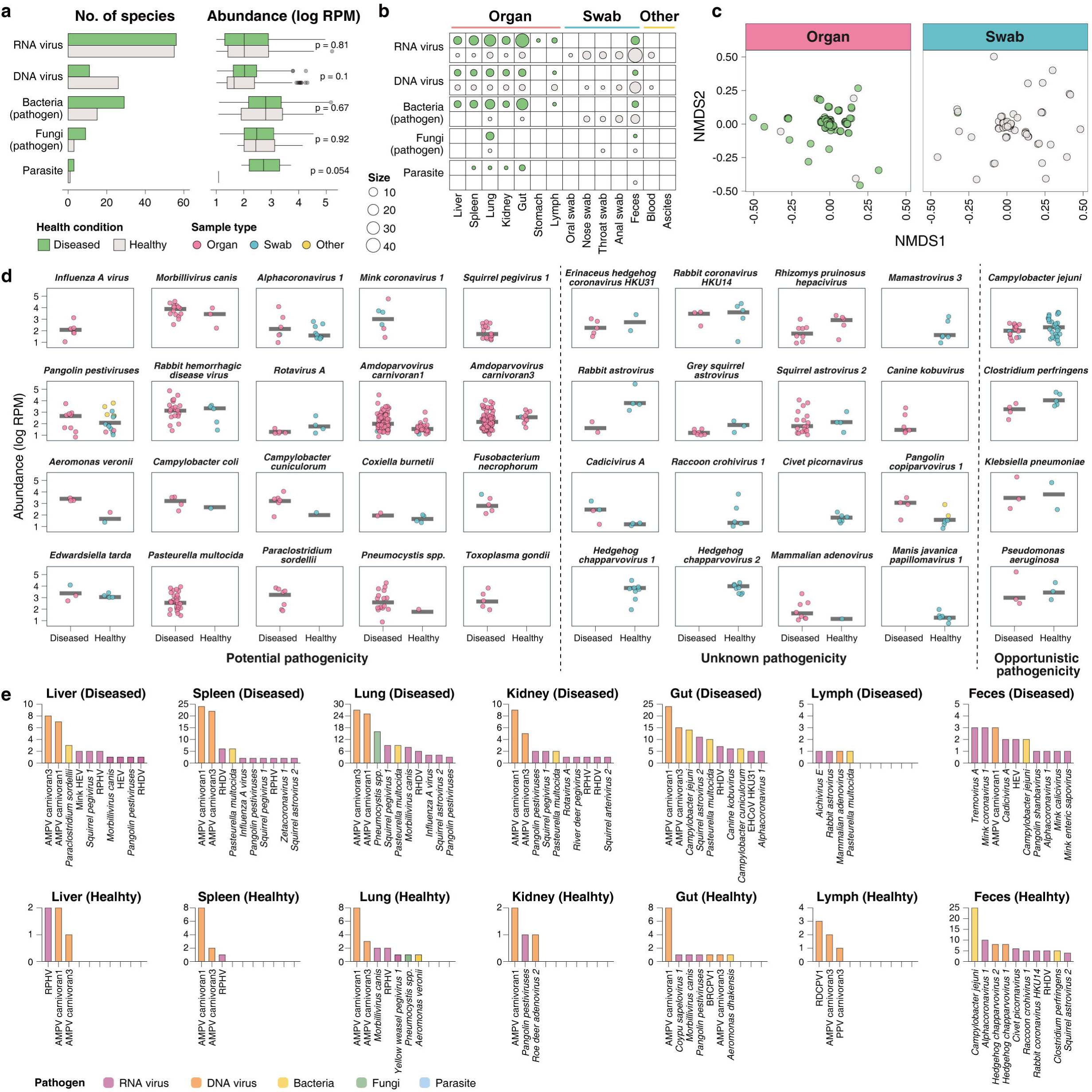
Comparisons of infectomes in diseased and healthy wild animals. a, Richness and abundance of each pathogen group are shown between diseased and healthy groups. The text in each pair box indicts the significance of abundance comparisons between diseased and healthy animals. **b,** Number of viral, bacterial, fungal and parasitic pathogens identified at the sample type level. The circle size reflects the number of pathogens detected. **c,** Non-metric multidimensional scaling (NMDS) analysis of a Bray-Curtis dissimilarity matrix calculated from organ and swab samples showing differences in pathogen distribution patterns between diseased and healthy animals. **d,** Abundance comparisons between healthy and unhealthy animals for pathogens detected at least five libraries (i.e., prevalence ≥5 libraries). **e,** Prevalence of pathogens in diseased and healthy tissues, and feces.

**Fig. 4.**
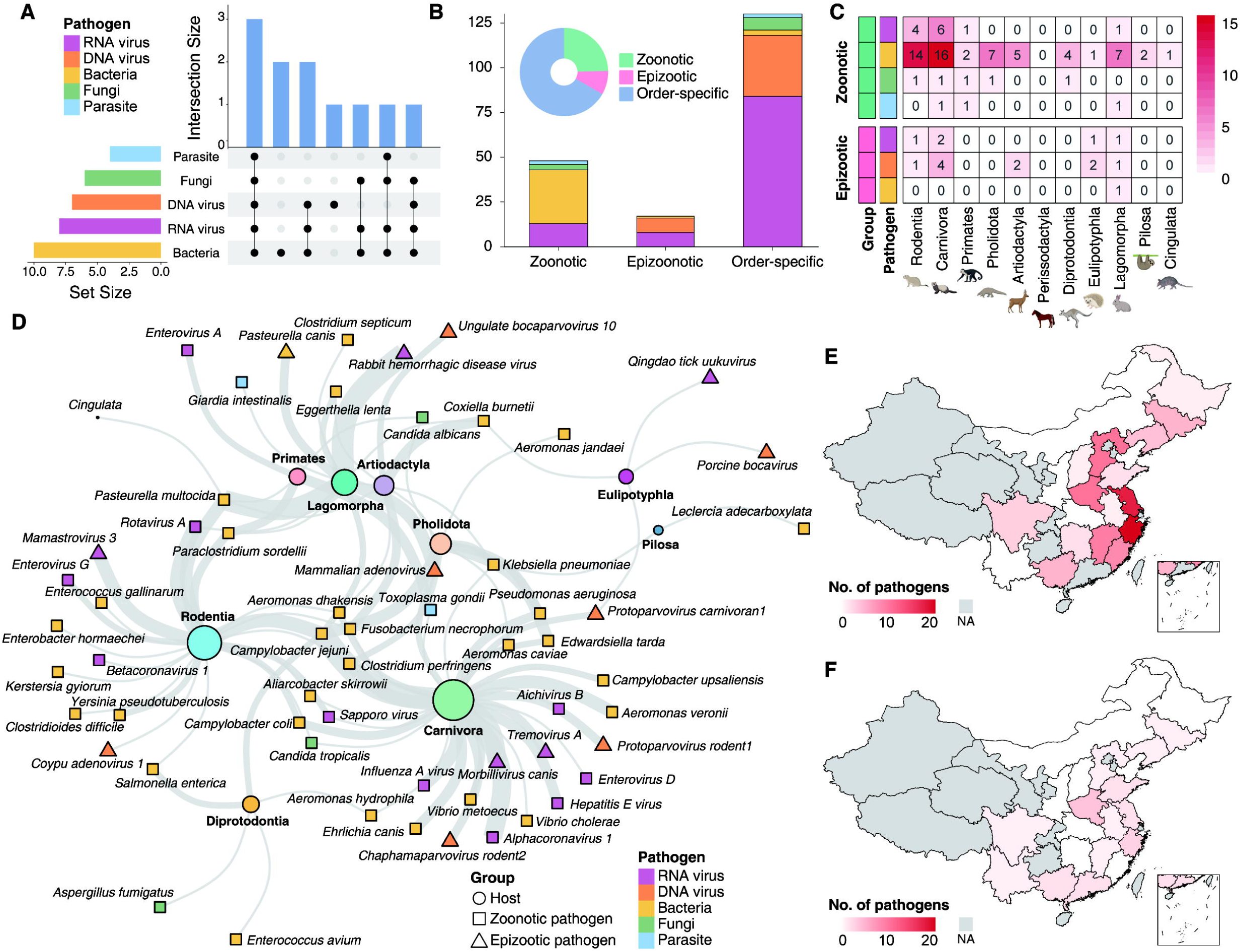
Epidemiological patterns of zoonotic and epizootic pathogens. a, The upset diagram illustrates the incidence of pathogen infection in mammalian orders. The left bar plot shows the number of infected animal orders of each pathogen group. The top right bar plot and crosspoint diagram shows the co-infection of pathogen groups in different animal orders. **b,** Number of (i) zoonotic (i.e., capable of transmission between animals and humans), (ii) epizootic (i.e., capable of transmission between animals from different orders), and (iii) order-specific (i.e., exclusively infecting mammalian species within the same order) pathogens identified in this study. **c,** The number of zoonotic and epizootic pathogens at the level of mammalian order. **d,** Associations between potential zoonotic and epizootic pathogens and mammalian hosts. The size of the colored circles representing animal hosts indicates the abundance of potential zoonotic and epizootic pathogens identified within each mammalian order, while the thickness of the line reflects the prevalence of each pathogen detected in that particular mammalian order. **e, f,** The number of zoonotic (**e**) and epizootic (**f**) pathogens at provincial level. “NA” shaped in grey denotes provinces for which the sample is not available.

In addition to viral pathogens, we identified a diverse array of bacterial, fungal, and parasitic pathogens, many of which are well-documented for their association with human diseases and zoonotic transmission. For instance, *Clostridioides difficile* is a leading cause of healthcare-associated infections^20^, while *Coxiella burnetii*, the causative agent of Q fever, poses significant zoonotic risks^21^. Similarly, *Pasteurella canis* which is commonly linked to animal bites^22^, a relative of *Aeromonas caviae* usually associated with gastrointestinal infections^23^, and the presence of parasites such as *Giardia intestinalis* and *Toxoplasma gondii*, underscore the broad spectrum of pathogens transmitted from animal reservoirs to humans. Additionally, we detected several opportunistic pathogens, including *Klebsiella pneumoniae*, *Pseudomonas aeruginosa*, *Campylobacter jejuni*, *Eggerthella lenta*, and *Aspergillus fumigatus*, that are generally associated with mild or opportunistic infections in immunocompromised individuals^24^. Notably, we also identified four divergent members of the fungal genus *Pneumocystis* which is associated with pneumonia in humans and mammalian hosts^7^. These fungi were primarily identified in species within the orders Carnivora and Rodentia (Fig. 1 and Extend Data Fig 5b). Together, these findings highlight the intricate interplay between wildlife, domestic animals, and human health, emphasizing the critical need for ongoing surveillance and proactive measures to monitor and mitigate zoonotic transmission risks.

### A novel segmented coronavirus in diseased lesser pandas

Of particular note, we discovered a novel, segmented coronavirus in lesser (red) pandas (*Ailurus fulgens*) that exhibited signs of disease. Phylogenetic analysis using the RNA-dependent RNA polymerase (RdRP) gene placed this virus, provisionally denoted Zetacoronavirus 1, within the family *Coronaviridae*, but distinct from other known coronaviruses genera, including the putative *Epsiloncoronavirus* found in intestines and lungs of *Meles* (i.e. European badgers)^25^ (Fig. 2a). Indeed, genomic analysis showed that this virus had less than 50% similarity to known coronaviruses (Extend Data Fig 6). Importantly, this virus was primarily found in the lung and spleen tissues of lesser pandas in Jiangsu and Fujian provinces, with a high viral load in the lungs (RPM 7,396.26, Fig. 2b), suggesting a link to respiratory illness and high mortality. Despite efforts, we did not obtain the unsegmented genome typical of other coronaviruses. Rather, the virus appears to be segmented, consisting of at least two segments: one with ORF1ab and RdRP (1.47 kb, PQ685031) and another with the spike and nucleocapsid proteins (7.66 kb, PQ685032) (Fig. 2c). Rapid amplification of cDNA ends (RACE) confirmed this genome structure and revealed a highly conserved 700 bp sequence at the 5’ terminus of both segments, with at least 11 SNPs between strains (Fig. 2d, Extend Data Fig 7). This structure resembles bi-segmented Nido- like viruses found in aquatic vertebrates, which exhibit an identical trailing sequence of 129 nucleotides at their respective 3’ ends (Fig. 2a and 2c)^26^. Both contigs exhibited poly(A) tails at their 3’ termini, confirmed through 3’ RACE, further supporting the presence of a bi-segmented genome (Fig. 2d, Extend Data Fig S7). This conserved 700 bp sequence at the 5’ terminus (5UTR-F-1: GGCAAGCAGTGGTATCAACGCA; 5UTR-F-2: ACCAACAGGCTTGACCTCTTGA) and poly(A) tails at the 3’ terminus were utilized in PCR attempts to amplify additional segments. However, no products were obtained, providing further evidence for the bi-segmented nature of the genome. We also utilized long-read amplicon sequencing data to validate the contiguity within both segments, especially the regions between the S and N genes, of Zetacoronavirus 1 (Extend Data Fig 7c).

Evolutionary analysis of RdRP and spike proteins revealed substantial genetic divergence of Zetacoronavirus 1 from known genera within the *Coronaviridae* (pairwise distances: RdRP > 0.781, spike > 1.025; Fig. 2e, Extend Data Fig 6b). We therefore proposed that this virus should be classified as a putative new genus, tentatively named Zetacoronavirus, based on its significant divergence and distinct genomic features. The virus segments, although small, contain key components for replication and structural (i.e., spike and nucleoprotein) functions. Detailed genome annotation revealed 13 non-structural proteins (NSPs) and an unconventional ribosomal frameshifting signal (TTTAAAT, instead of the canonical TTTAAAC) within segment 1 (PQ685031), encoding ORF1ab. Interestingly, three non-structural proteins – NSP1, NSP2, and NSP15 – were absent. Segment 2 (PQ685032), encoding the spike and nucleocapsid proteins, notably lacked the envelope and membrane structural proteins. These distinct features define a compact, bi-segmented genome architecture (Fig. 2), highlighting the hidden diversity of mammalian coronaviruses and their largely unexplored genomic formations. These observations merit additional virus isolation and molecular validation.

### Comparisons of infectomes in diseased and healthy wild animals

To assess disease associations of the pathogens identified here, we compared infectomes from diseased and healthy wild animals. The diseased group shown a higher incidence of RNA viruses, bacteria, fungi, and parasites, while DNA viruses were more common in the healthy group, suggesting opportunistic or less-pathogenic roles (Fig. 3a). However, differences in pathogen distribution were partly influenced by sample type: organ samples predominated in the diseased group, while swab samples were more common in the healthy group (Figs. 3b and 3c). Several pathogens, including *Influenza A virus*, *Morbillivirus canis*, *Alphacoronavirus 1*, were more abundant in diseased animals, indicating a potential link to illness (Fig. 3d). Conversely, pathogens like *Campylobacter jejuni* and *Klebsiella pneumoniae* were equally prevalent in both groups, suggesting opportunistic or subclinical roles (Fig. 3d).

We also examined tissue tropisms, which revealed greater pathogen richness in the gut and lungs compared to other organs such as the liver, spleen and kidney (Extend Data Fig 8a and 8b). Some pathogens, like AMPV carnivoran1 and RHDV detected in mink and rabbits, respectively, were present in multiple organs, suggesting systemic infections (Fig. 3e and Extend Data Fig 8c). Other pathogens showed the expected patterns of organ specificity; for example, gastrointestinal pathogens were mainly found in the gut, while respiratory pathogens, including newly identified *Pneumocystis* species, were found only in lung tissues (Fig. 3e and Extend Data Fig 8d). Overall, while many pathogens were distributed across multiple organs, most exhibited pronounced organ specificity. The novel Zetacoronavirus 1 appears to initially affect the lungs, although with evidence of spread to the spleen, raising concerns about its impact on multiple organs and systems.

Although healthy controls were not included, we identified key pathogens in several cases that may be associated with the observed diseases. For example, a ring-tailed lemur (*Lemur catta*) from Zhejiang province, which died in a zoo, exhibited necrotic liver lesions, hemorrhagic infarctions in a blackened and hardened spleen, severe pulmonary haemorrhage, and enlarged kidneys upon necropsy. A high load of *Toxoplasma gondii* was detected in various tissues, with RPM values ranging from 82 to 5,245 suggesting it was the likely causative agent. In five mink (*Neovison vison*) from Liaoning Province, symptoms of emaciation and yellow-green mucoid diarrhea were observed, and fecal samples revealed multiple pathogens linked to gastrointestinal diseases, including *Canine norovirus 2*, *Mink enteric sapovirus*, *Mink coronavirus 1*, *Amdoparvovirus carnivoran 1*, and *Campylobacter jejuni*. Additionally, continuous monitoring from 2021 to 2022 of a single Malayan pangolin (*Manis javanica*) revealed increasing viral loads of *Pangolin pestivirus 4*, *Pangolin shanbavirus*, and *Pangolin copiparvovirus 1* in a deceased individual. These viral infections likely contributed to the animal’s death, potentially linked to observed symptoms such as pericardial effusion, yellow discoloration, bladder distension, and ascites. Moreover, a tick-borne virus, Qingdao tick uukuvirus ^27^, was detected in hedgehogs with lung tissue calcification.

### Characterization of zoonotic and epizootic pathogens

We analysed the data generated here combined with all publicly available data to examine the frequency of host jumps among zoonotic and epizootic pathogens. These results revealed that species from the mammalian orders Carnivora, Rodentia, and Lagomorpha commonly host RNA viruses, DNA viruses, bacteria, fungi, and parasites (Fig. 4a). We then categorized the total set of 195 pathogens into three types: zoonotic (i.e., transmitting between animals and humans), epizootic (spreading across different animal orders), and order-specific (infecting species within the same order) (Fig. 4b and Supplementary Table 3). In this manner we identified 48 zoonotic, 17 epizootic and 130 order-specific pathogens, with Carnivora and Rodentia serving as both sources and recipients of zoonotic spillovers (Fig. 4c and Extend Data Fig 9). Pathogens at high prevalence and that exhibited evidence of cross-species transmission were detected in multiple animal species across different orders (Fig. 4d). Examples include the highly pathogenic avian influenza virus (HPAIV) H5N1 and H5N6 in raccoon dogs and mink, respectively, *Qingdao tick uukuvirus* in hedgehogs, and *Morbillivirus canis* (*Canine distemper virus*, CDV) in raccoon dogs, civets, mink, and other carnivores, highlighting their potential for cross-species transmission.

Phylogenetic analysis of the HA gene revealed that the H5N1 influenza virus identified in raccoon dogs from Shandong province was closely related to strains detected in chickens from Tottori and Chiba, all circulating in 2022. Similarly, the H5N6 influenza virus detected in mink from Shandong clustered with a human-associated strain identified in 2021 (Extend Data Fig 10). These findings raise concern, particularly as H5N1 has caused the deaths of thousands of marine mammals in South America since 2022 ^28,29^. We also identified known zoonotic pathogens such as *Sapporo virus*, *Campylobacter spp*., *Clostridium perfringens*, *Fusobacterium necrophorum*, *Toxoplasma gondii*, and *Coxiella burnetii* across multiple mammalian hosts, particularly in Zhejiang and Jiangsu provinces in southern China where nearly 20 cross-species transmissions were recorded here (Fig. 4e and 4f).

In addition to zoonotic pathogens, the data generated here highlight the importance of monitoring pathogens that are currently confined to individual animal populations. For example, despite their limited host range, pathogens like *Amdoparvovirus carnivoran1* (*Aleutian mink disease parvovirus*, AMDV), *Amdoparvovirus carnivoran3* (raccoon dog and arctic fox amdoparvovirus, RDAV), and pangolin pestiviruses could pose significant threats to the principal host as they are commonly associated with overt disease (Extend Data Fig 11).

## Discussion

Wild animals are often linked to the emergence of major human diseases, making it crucial to identify existing or potential pathogens in these species. Such efforts can also help trace the origins of specific epidemics and assess the risk of future outbreaks, particularly the identification of host “generalist” pathogens that seemingly possess an enhanced ability to emerge in new species, including those from different mammalian orders^4,15,16,30^. The diverse array of pathogens uncovered in this study highlights the importance of adopting a comprehensive infectome approach to better understand and mitigate the risks of zoonotic disease transmission ^18,19^. By profiling the spectrum of viral, bacterial, fungal, and parasitic pathogens we have demonstrated the intricate and interconnected nature of wildlife-pathogen dynamics. Traditional pathogen-specific or virus-focused studies risk overlooking important contributors to disease ecology, including bacterial, fungal, or parasitic agents capable of zoonotic or epizootic transmission. For example, the identification of opportunistic pathogens such as *Pseudomonas aeruginosa* and *Aspergillus fumigatus* underscores their potential roles in wildlife health and spillover events. As such, these findings emphasize the need for integrative surveillance that encompasses all pathogen types, enabling more accurate risk assessments and informed, proactive interventions.

The complex interplay between wildlife pathogens and human health underscores the need for a collaborative, interdisciplinary approach to disease surveillance. By including diverse mammalian species, particularly under-sampled orders like Carnivora and Rodentia, this study highlights the critical importance of understanding pathogen reservoirs in both wild and managed animal populations. For instance, the high pathogen loads observed in fur-farmed carnivores suggest that anthropogenic pressures may significantly amplify spillover risks^31^. Additionally, the discovery of novel pathogens such as Zetacoronavirus 1 in lesser pandas reinforces the role of wildlife as reservoirs for emerging infectious diseases. These findings call for enhanced collaboration among ecologists, veterinarians, epidemiologists, and public health experts to develop comprehensive frameworks for pathogen monitoring. Such frameworks should integrate genomic, ecological, and epidemiological data to better predict and mitigate future outbreaks.

## Limitations

A key limitation of this study stems from differences in sample types collected from diseased and healthy groups, which may have influenced the observed variations in pathogen diversity and abundance. In particular, diseased samples primarily consisted of internal organ tissues, such as the liver, spleen, lungs, and intestines—sites that are more likely to harbor pathogens directly associated with clinical symptoms. In contrast, healthy samples were largely derived from fecal or respiratory swabs, which may capture a different subset of the infectome. These methodological discrepancies complicate the interpretation of whether the observed differences in pathogen richness and load between the groups are truly disease-associated or artifacts of sample type variability. Additionally, this study is geographically limited to China, highlighting the need to expand the framework to encompass a wider range of species and geographic regions for more comprehensive global health preparedness. Furthermore, while we relied on genomic tools for pathogen characterization, the isolation and functional study of novel agents in cell and animal models will be crucial for gaining deeper insights into their potential impacts on wildlife and human populations.

## Methods

### Sample collection

Wild animal sampling was conducted from 2019 to 2024 across natural habitats, artificial breeding sites, and zoos in 20 Chinese provinces (table S1). A total of 1,922 samples were collected from 57 mammalian species, spanning 11 orders and including 1,283 samples from diseased animals and 639 from healthy controls. The animal host examined here included 9 species from Rodentia (*n*=315), 19 from Carnivora (*n*=745), 13 from Primates (*n*=78), two from Pholidota (*n*=171), 14 from Artiodactyla (*n*=237), three from Perissodactyla (*n*=16), three species Diprotodontia (*n*=24), one from Eulipotyphla (Amur hedgehog, *n*=100), and one species each from Lagomorpha (*n*=229), Pilosa (*n*=1) and Cingulata (*n*=6) (Fig. 1A and fig. S1). Tissue samples from organs such as the liver, spleen, lung, kidney, intestine, stomach, and lymph were primarily obtained from deceased animals, whereas oral, nasal, throat, anal and stool swabs were mainly taken from healthy animals. All samples were transported on dry ice and stored in a -80 freezer.

Host species identification was initially conducted by experienced field biologists based on morphological characteristics at the time of capture. This identification was further validated through cytochrome c oxidase subunit I (*COX1*) gene analysis using RNA sequenceing data generatied for each library.

The sampling and sample processing procedures were approved by the Ethics Committee of Sun Yat-sen University (SYSU-IACUC-MED-2021-B0123).

### Sample processing and sequencing

RNA extraction and sequencing were performed for each sample. Total RNA was extracted from organ tissues using RNeasy Plus mini kit (QIAGEN, Germany) and from swabs and blood using the RNeasy Plus universal mini Kit (QIAGEN, Germany), following the manufacturer’s protocols. Meta-transcriptomic libraries for each pool were prepared using MGIEasy RNA library prep Kit V3.0. The resulting libraries were sequenced on the DNBSEQ T series platform (MGI), generating 150bp paired-end meta-transcriptomic reads.

### Pathogen identification and confirmation

For each sequence data set, the majority of ribosomal RNA (rRNA) reads were initially removed using URMAP (version 1.0.1480)^32^. Adapters, duplicate and low-quality reads were filtered out using fastp (version 0.20.1, parameters: -q 20, -n 5,-l 50,-y, -c, -D)^33^. The reads with low complexity were removed using PRINSEQ++ (version 1.2, options: -lc_entropy=0.5 - lc_dust=0.5)^34^. Residual rRNA reads were further eliminated by mapping to the SILVA rRNA database (Release 138.1) 70 using Bowtie2^35^. Unless otherwise specified, all software was run with default settings. This step removed low-quality reads, adaptor sequences, non-complex reads, and duplicates. The remaining reads were then assembled *de novo* using MEGAHIT program (version 1.2.9)^36^ with default parameters. The resulting contigs were annotated by comparing them against the NCBI non-redundant (nr) protein database (version 2023.01.09) using Diamond blastx (version v2.1.9.163)^37^, with an *E*-value threshold of 1*E*-3 to balance high sensitivity with low false-positive rate.

To identify viral pathogens, taxonomic lineage information for each contig’s top blast hit was retrieved, with those classified under the kingdom “Viruses” identified as potential viral hits. Contigs with unassembled overlaps were merged using the SeqMan program implemented in the Lasergene software package (DNAstar, version 7.1)^38^. Final virus genomes were validated by mapping reads to the corresponding contigs and examining the mapping results using the Geneious software package^39^. Viral contigs likely associated with vertebrates, particularly vertebrate-specific and vector-borne vertebrate viruses, were initially identified based on the taxonomic lineage from blastx results and confirmed through phylogenetic analyses. Contigs clustering within a vertebrate-specific or vector-borne virus family/genus/group were further classified as vertebrate viruses. Viral sequences/genomes showing <80% nucleotide similarity identity to known viruses were assigned as potential novel virus species, which were further validated via phylogenetic analysis.

To identify bacterial, fungal and parasitic pathogens, we first employed MetaPhlAn (version 4.0.0)^40^ to detect potential pathogens within these groups. Relevant background marker genes for bacteria, fungi and parasites were downloaded from GenBank and subjected to blastn comparison^41^ (version 2.15.0+) to obtain contigs that contain these marker genes. These contigs were then further compared against the nt database (version 2023.04.23) to determine microbial taxonomy at the species level. The identified pathogen species were confirmed through phylogenetic analysis of specific marker genes, such as rpoB for bacteria, COI for fungi, GP60, GDH and COI for parasite.

### Estimation of prevalence and abundance

To estimate the relative abundance of pathogens in each library, quality-trimmed reads were first aligned to the SILVA database (www.arb-silva.de, version 132.1) using Bowtie2 (version 2.5.2)^35^ to remove ribosomal RNA reads. Unmapped reads were then aligned to confirmed viral genomes using the “end-to-end” setting, and the abundance of each virus species was determined using the following formula: total viral reads divided by total non-redundant reads, multiplied by 1,000,000 (i.e., mapped reads per million total reads [RPM]). For bacterial, fungal and parasitic pathogens, abundance was also calculated as RPM based on reads mapped to relevant bacterial fungal and parasitic genomes downloaded from GenBank.

A pathogen was considered “positive” based on two established criteria. First, to mitigate false-positives from index hopping (a common issue in high-throughput sequencing where reads from one sample are incorrectly assigned to another), we set a read abundance threshold relative to the highest read count in each sequencing lane. If the total read count of a specific pathogen in a library was less than 0.2% of the highest read count for that pathogen within the same sequencing lane, it was considered a false positive due to index-hopping. Additionally, pathogens with fewer than 10 RPM were classified as “negative” and excluded from further statistical analysis, as they were likely false positives. These thresholds were validated in previous publications through reverse transcription polymerase chain reaction (RT-PCR) re- confirmation^19,42^.

### RACE and long-read sequencing for segmented Zetacoronavirus 1

Total RNA was used for 5’ and 3’ RACE (Rapid Amplification of cDNA Ends) amplification to capture the full-length sequences of the S and ORF1ab genes of a segmented coronavirus identified in the lesser panda. The amplification was performed using the HiScript-TS 5’/3’ RACE Kit (Vazyme, Nanjing, China), following the manufacturer’s protocol. First, the first-strand cDNA was synthesized from the total RNA using the RT enzyme and the CDS Primer.

Gene-specific primers were designed to amplify the 5’ and 3’ ends of the S and ORF1ab genes. PCR amplification was carried out under the following conditions: 30 cycles of denaturation at 98°C for 10 seconds, annealing at 68°C for 15 seconds, and extension at 72°C for 3 minutes, with a final extension at 72°C for 5 minutes. The amplified PCR products were cloned into the pClone007 vector using the pClone007 Blunt Simple Vector Kit (Tsingke, Beijing, China). The recombinant plasmids were verified by Sanger sequencing, which was conducted by Sangon Biotech (Shanghai, China). The sequences of the primers used were as follows: S-3’ RACE forward: ACTGTTGTGTATGTTCTTGAGAAC, S-5’ RACE reverse: GCATACGGTACAATGGACTTA, ORF1ab-3’ RACE forward: CGTGGCTAATGACATTAGACCAT, ORF1ab-5’ RACE reverse: CTCGAGCTTAACACAACCCGTAATT.

A 22-plex ATOPlex panel, targeting 1,500-bp amplicons, was designed to comprehensively cover the genome of Zetacoronavirus 1 (https://atoplex.mgi-tech.com). RNA was reverse transcribed into complementary DNA (cDNA) using random hexamer primers, and whole-genome amplification was subsequently performed with an equimolar mixture of 22-plex primers, following the manufacturer’s protocol (MGI, Shenzhen, China). The amplified products were then subjected to end repair and adapter ligation with CycloneSEQ adapter, followed by long-read sequencing on the CycloneSEQ platform (MGI, Shenzhen, China).

### Evolutionary analysis

We conducted phylogenetic analysis to verify that the viruses were vertebrate-associated viruses. Initially, these viral sequences were classified into their respective viral family or genera based on the Diamond blastx results. Subsequently, the amino acid sequences of conserved marker genes (i.e., RdRP for RNA viruses and capsid protein for DNA viruses) from these virus genomes were aligned with related viral reference sequences downloaded from GenBank using MAFFT (version 7.475)^43^, employing the L-INS-I algorithm. The reference sequences, selected to represent diversity and the top hit for each virus species identified here, were incorporated into each family. Any ambiguous regions were removed using TrimAL (version 1.4.rev15)^44^.

Phylogenetic trees were inferred for the sequence alignment from each virus family using the maximum likelihood method in PhyML (version 3.1)^45^, employing the LG model of amino acid substitution and a subtree pruning and regrafting (SPR) branch swapping algorithm. A similar approach was applied for the phylogenetic analysis of bacteria, fungi, and parasites. However, due to high sequence identity, genome-scale nucleotide sequences were used in these cases, and a general-time reversible (GTR) substitution model was employed for phylogenetic inference.

### Genome annotation of a newly identified coronavirus

The annotation of potential open reading frames (ORFs) in coronavirus genomes, as shown in Fig. 2, was predicted based on two main criteria: (i) the predicted amino acid sequences were longer than 100 amino acids in length, and (ii) they were not fully nested within larger ORFs. The ORF annotation process primarily relied on comparisons of predicted proteins to hidden Markov models (HMMs) from the Pfam database (version 2022.03.23, https://pfam-legacy.xfam.org/) using hmmscan v3.3.2 (implemented in HMMER, with parameters *e*=1*E*-3 and score ≥30)^46^. Additionally, Diamond blastp (version v2.1.9.163)^37^ was used for further annotation, comparing sequences against the nr protein database (version 2023.01.09) with an e-value threshold of 1*E*-3.

### Definition of zoonotic and epizootic pathogens

To assess the risk of cross-species transmission for all pathogens in this study, we compiled host information for each specific pathogen by integrating data from the NCBI nucleotide database with the host information of pathogens identified from animal samples collected in this study.

Based on their host range, the 195 pathogens were categorized into three groups: (i) zoonotic pathogens (capable of transmission between animals and humans), (ii) epizootic pathogens (capable of transmission between animals from different orders), and (iii) order-specific pathogens (infecting animal species within the same order).

## Data availability

The genome sequences of bi-segmented coronavirus identified in lesser pandas are available at the NCBI GenBank database under accession numbers PQ685031-PQ685032.

## Supporting information

Supplementary tabe 1

Supplementary tabe 2

Supplementary tabe 3

## Acknowledgments

This work was supported by the National Key Research and Development Program of China (2024YFC2607502), the National Natural Science Foundation of China (82341118, 32270160), the Shenzhen Science and Technology Program (KQTD20200820145822023), the Natural Science Foundation of Guangdong Province (2022A1515011854), the Guangdong Province “Pearl River Talent Plan” Innovation and Entrepreneurship Team Project (2019ZT08Y464), the Hong Kong Innovation and Technology Fund (ITF) (MRP/071/20X), and the Health and Medical Research Fund (COVID190206). E.C.H. is funded by a National Health and Medical Research Council (Australia) Investigator grant (GNT2017197) and by AIR@InnoHK administered by the Innovation and Technology Commission, Hong Kong Special Administrative Region, China. We gratefully acknowledge colleagues at BGI Research and the China National Genebank (CNGB) for library preparation, sequencing and data management.

## Author contributions

Conceptualization: X.H., E.C.H., M.S.; Methodology: X.H., J.W., D.-X.W., E.C.H., Z.Q.D., M.S.; Investigation: X.H., J.W., D-X.W., W-C.W., S-J.L., S-Q.M., H-L.L., Y.F., L-F.Y., P-B.S., Z-R.R., L.Y., Z-Q.D.; Sampling: F-F.C., X-X.L., H-J.H., W-J.S., J.C., M-M.Z., Z-W.J., X-M.L.; Visualization: X.H.; Funding acquisition: D.Y., E.C.H., M.S.; Writing – original draft: X.H., E.C.H., M.S.; Writing – review & editing: All authors; Supervision: X.H., Z.Q.D., M.S.

---

**Extended Data** Fig. 1**. Sample size of each mammalian species, including sample type, sample location and health condition.** The left heatmap displays the sample size of each species at the level of sample type, including organ tissues and swabs, with the top bar plot marking the sample size for each specific sample type. The middle heatmap displays the sample size of each species at the sample location level, with the top bar plot marking the sample size for each specific sample location. The right heatmap displays the sample size of each species at the health condition level, with the top bar plot marking the sample size for healthy and diseased groups. The rightmost bar plots display the sample size of each animal species. The gradation of red color reflects the sample size. Animal species are colored according to their animal order.

**Extended Data** Fig. 2**. Number of potential pathogens detected in each mammalian order.** The bar plots illustrate the number of potential pathogens detected in each viral family or non-viral genus for every mammalian order. Viral family or non-viral genus is colored according to their pathogen group.

**Extended Data** Fig. 3**. Abundance of potential pathogens detected in each mammalian order.** The box plots illustrate the abundance of potential pathogens detected in each viral family or non-viral genus for every mammalian order. Viral family or non-viral genus is colored according to their pathogen group.

**Extended Data** Fig. 4**. Distribution of potential pathogens detected in each mammalian species.** Heatmap showing the richness and abundance of infectome in wild mammals at the family and genus levels. Pathogen groups and animal orders are shown by the colors on the heatmap. The gradation of cell red color reflects the abundance.

**Extended Data** Fig. 5**. Total infectome in wild mammals. a,** Heatmap showing the richness and abundance of RNA virus in wild mammals. The newly discovered coronavirus in lesser pandas is highlighted with red font. **b,** Heatmap showing the richness and abundance of DNA virus, bacteria, fungi and parasites in wild mammals. Pathogen groups and mammalian orders are shown by the colors on the heatmap. The gradation of cell red color reflects the abundance.

**Extended Data** Fig. 6**. Genomic similarity comparison between Zetacoronavirus 1 and other known coronaviruses. a,** Blastx results of ORF1ab and spike protein. **b,** Pairwise distance results of ORF1ab and spike protein sequences.

**Extended Data** Fig. 7**. The Rapid Amplification of cDNA Ends (RACE) method and long-read sequencing for two segments of Zetacoronavirus 1. a,** Primers used for the RACE method. Specific primers designed for amplifying the 5’ and 3’ ends of the S gene and ORF1ab gene are listed, including their corresponding names and sequences. **b,** Agarose electrophoresis results of the RACE method. Lane 1: 2 kb DNA ladder; Lane 2: amplified product of the 3’ end of the S gene; Lane 3: amplified product of the 5’ end of the S gene; Lane 4: amplified product of the 3’ end of the ORF1ab gene; Lane 5: amplified product of the 5’ end of the ORF1ab gene. Clear bands indicate the successful amplification of the 5’ and 3’ ends of the S gene and ORF1ab gene. **c,** The long-read coverage plot of two segments of Zetacoronavirus 1. The coverage plot illustrates mapping depth of long reads across the two segments of Zetacoronavirus 1.

**Extended Data** Fig. 8**. Tissue tropism of potential pathogens detected in this study. a,** Number of potential pathogens detected in each organ tissue. **b,** Networking graph showing co-infection of pathogens between different organ tissues. **c,** Heatmap showing multi-organ infection of pathogens detected here. **d,** Heatmap showing distribution and abundance of pathogens with organ-specificity.

**Extended Data** Fig. 9**. Prevalence of zoonotic and epizootic pathogens in each mammalian species.** The gradation of the red color and number in cells reflects the prevalence. Pathogen groups and animal groups are shown by the colors on the heatmap. The right bar plots showing the number of zoonotic and epizootic pathogens detected in each mammalian species.

**Extended Data** Fig. 10**. Phylogenetic tree of the H5N1 and H5N6 subtypes of influenza A virus based on the hemagglutinin (HA) and neuraminidase (NA) genes.** Inter-specific tree showing the phylogenetic diversity of H5N1 and H5N6 identified in this study. All trees were midpoint-rooted for clarity only, and the scale bar indicates 0.05 nucleotide substitutions per site. The sequences identified here are marked by orange with circle. The strains from human, other mammals, avian and environment are indicated by red, orange, blue and green, respectively.

**Extended Data** Fig. 11**. Phylogenetic tree of order-specific pathogens.** Inter-specific tree showing the phylogenetic diversity of *Amdoparvovirus carnivoran1*, *Amdoparvovirus carnivoran3*, *Pangolin pesitivirus*, *Pangolin shanbavirus*, *Canine norovirus 2*, *Mammalian enterovirus* and *Pneumocystis spp* identified in this study. All trees were midpoint-rooted for clarity only, and the scale bar indicates 0.05 nucleotide substitutions per site. The sequences identified here are marked by colored dot.

**Supplementary Table 1.** Detailed information on all animal samples collected in this study.

**Supplementary Table 2.** The normalized abundance levels (measured by RPM) of each pathogen in different sequencing libraries.

**Supplementary Table 3.** Detailed information on the classification of zoonotic, epizootic and order-specific pathogens identified in this study.

