## Supplementary figures and images for "Comprehensive Infectome Analysis Reveals Diverse Infectious Agents with Zoonotic Potential in Wildlife"

### Extended Data Fig. 1

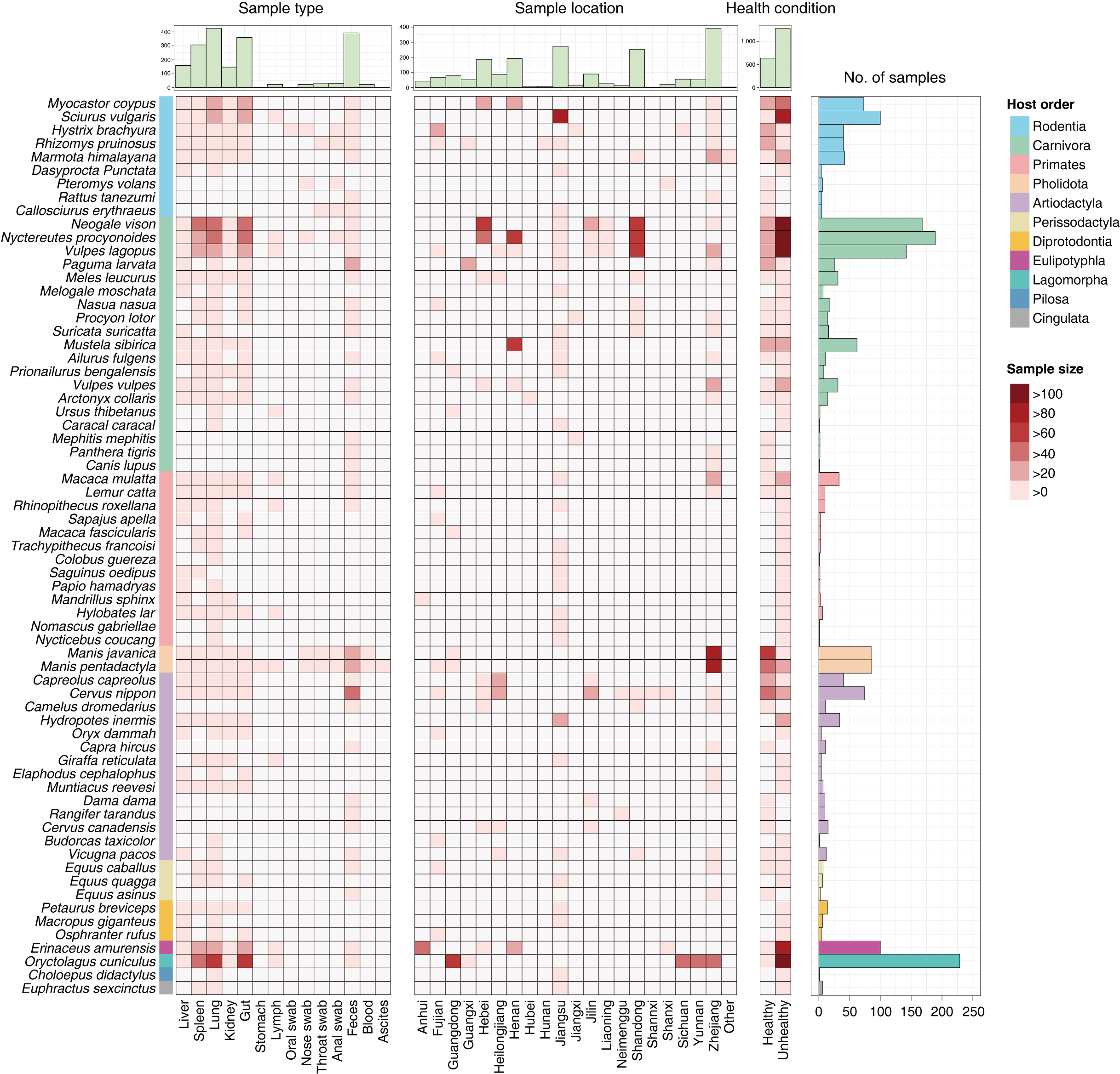

### Extended Data Fig. 2

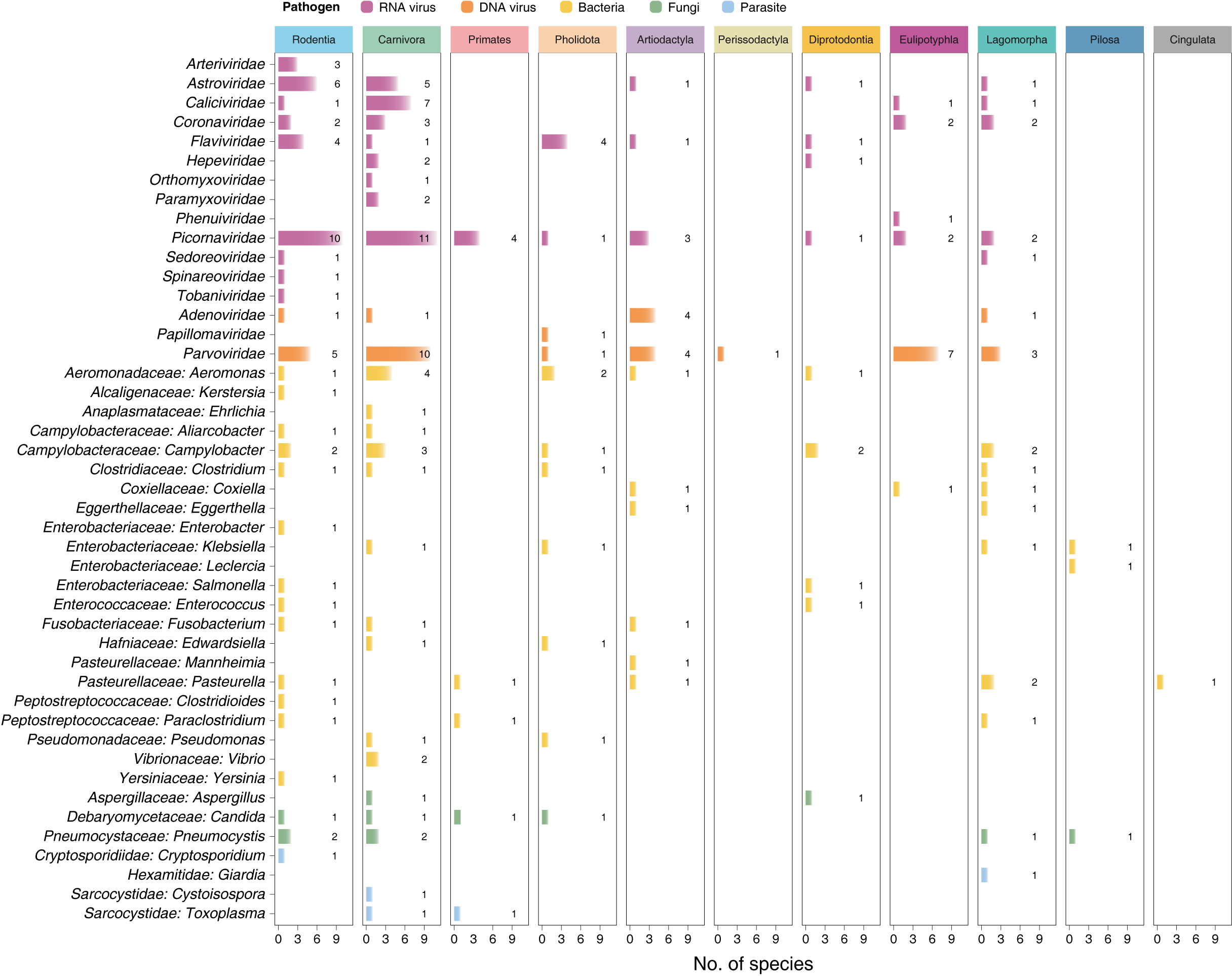

### Extended Data Fig. 3

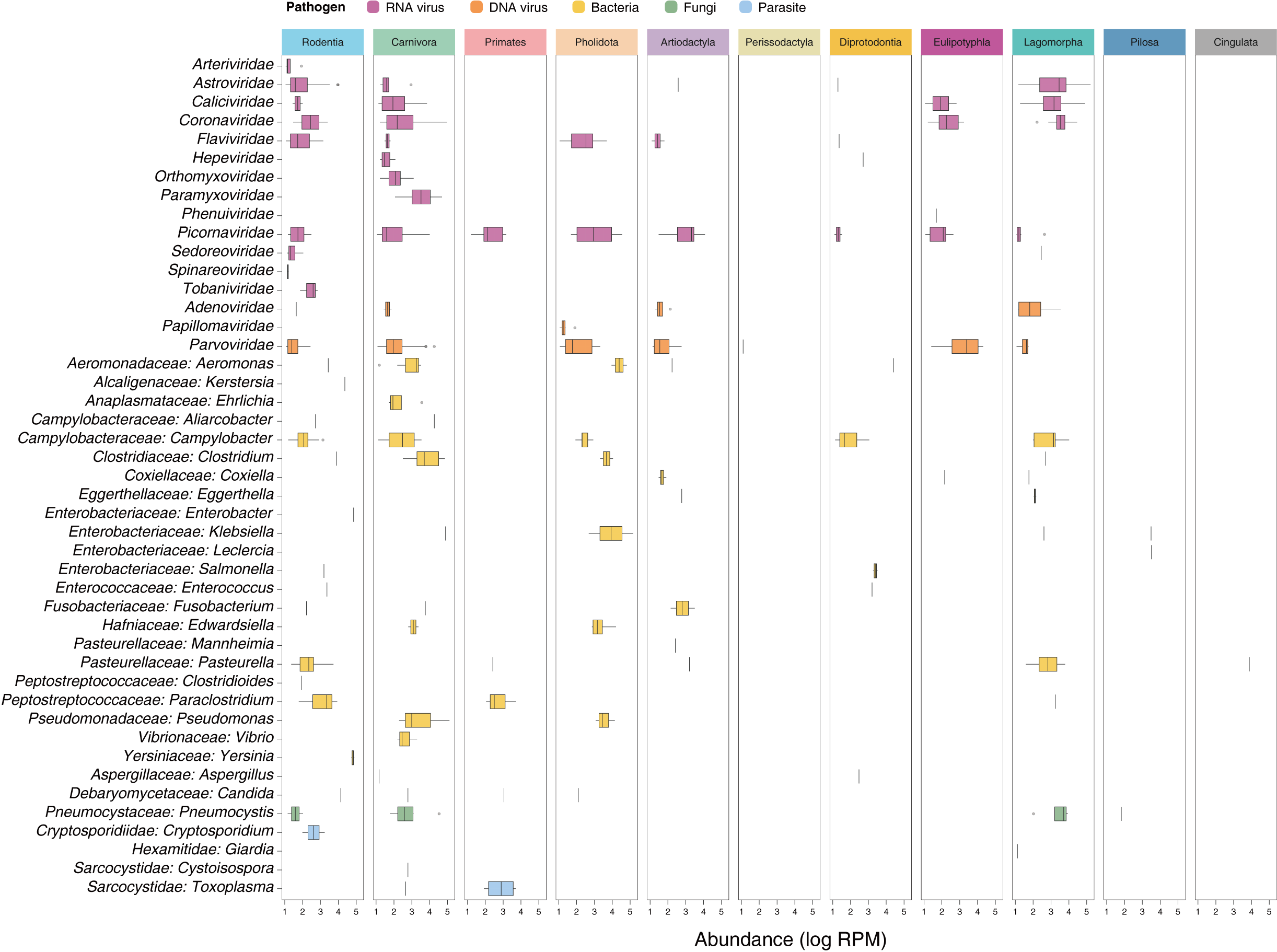

### Extended Data Fig. 4

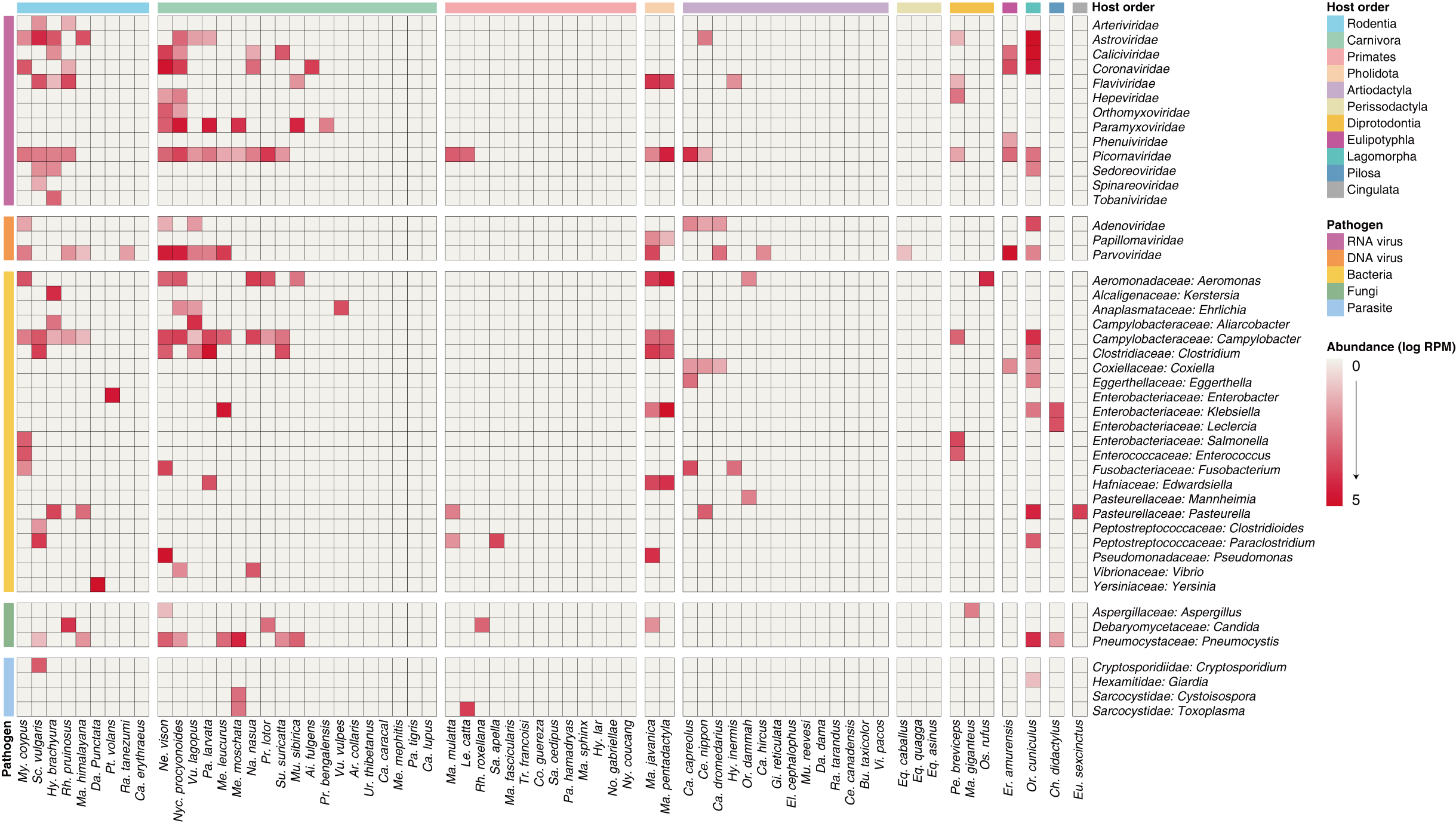

### Extended Data Fig. 5

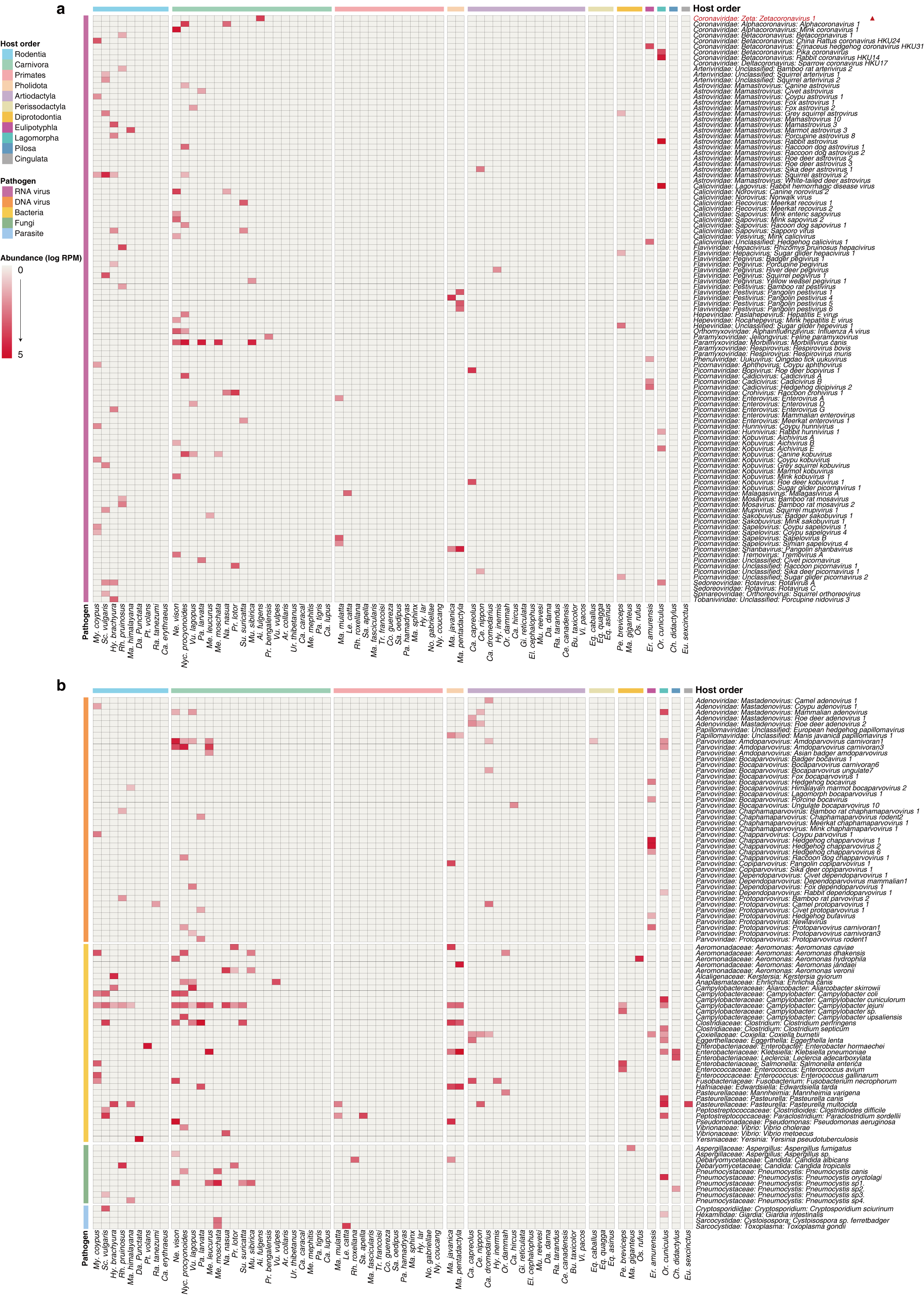

### Extended Data Fig. 6

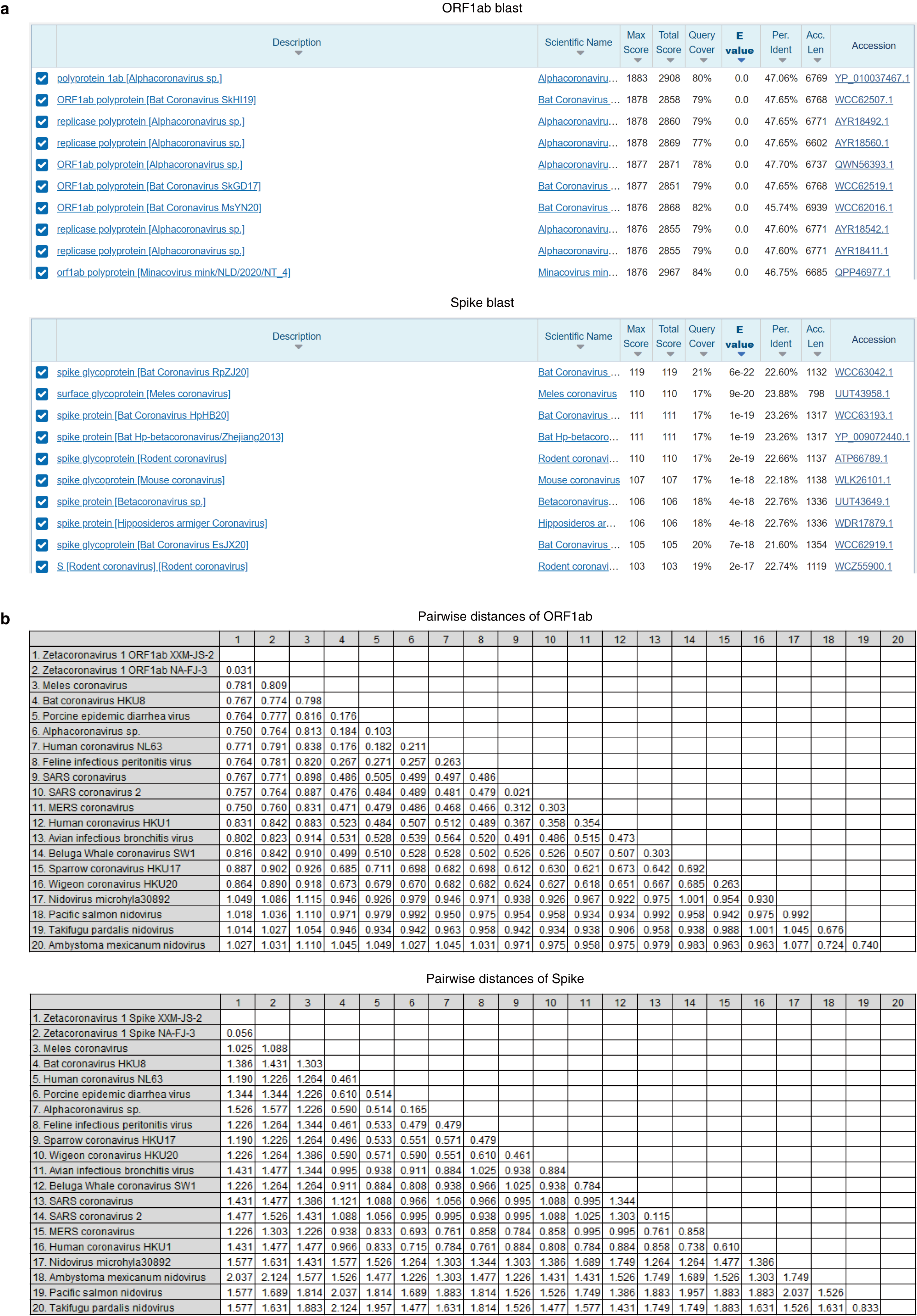

### Extended Data Fig. 7

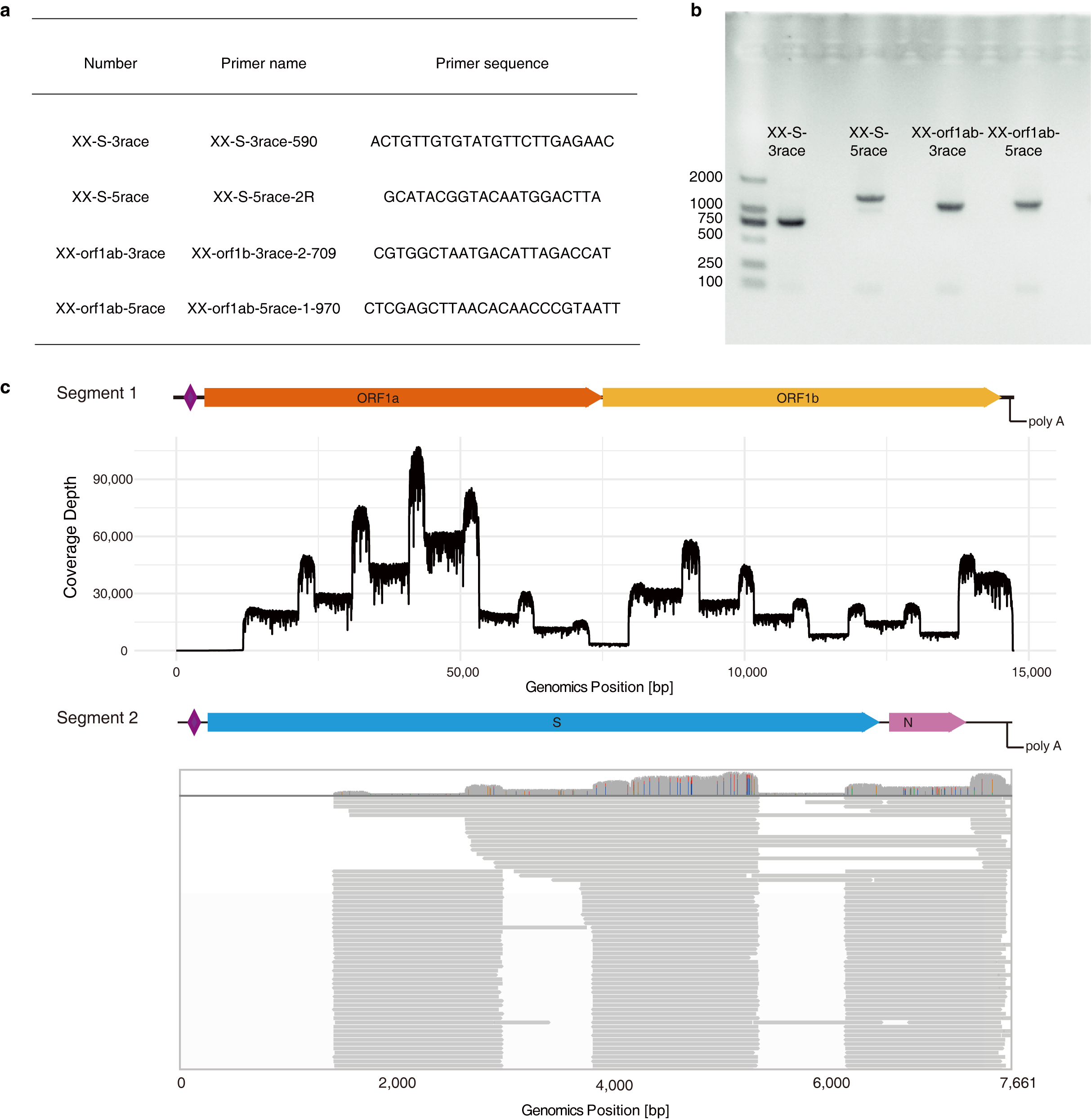

### Extended Data Fig. 8

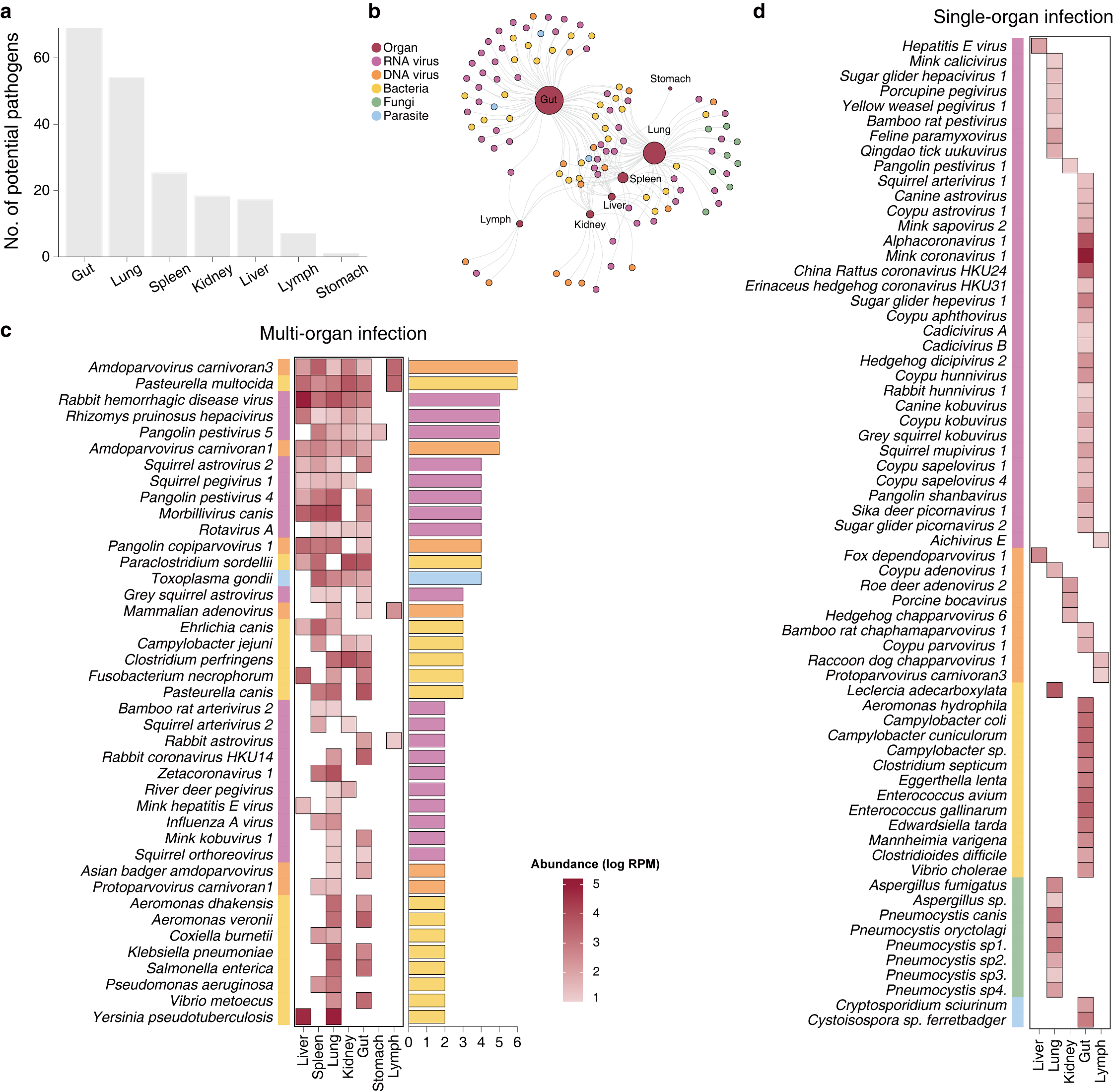

### Extended Data Fig. 9

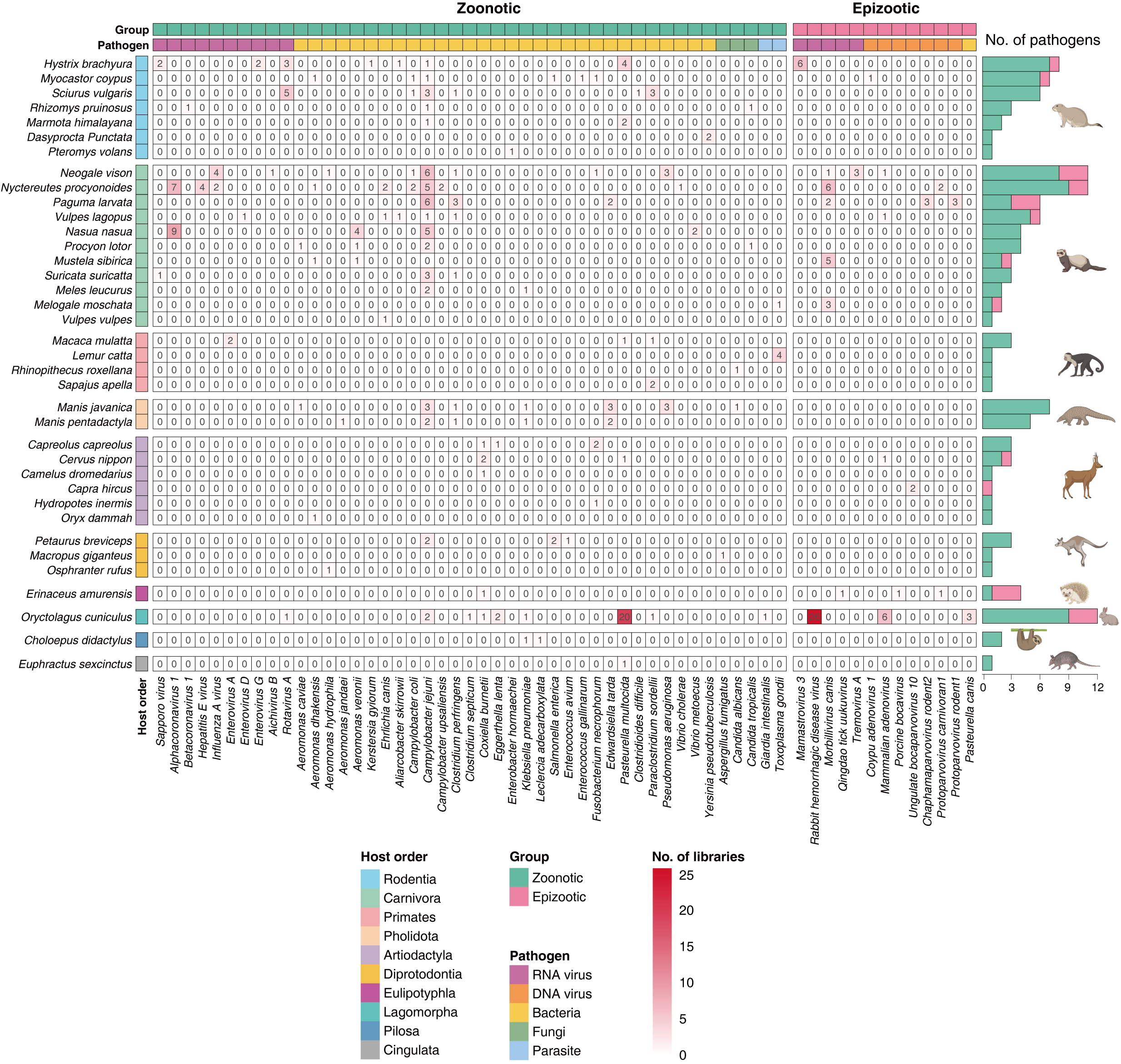

### Extended Data Fig. 10

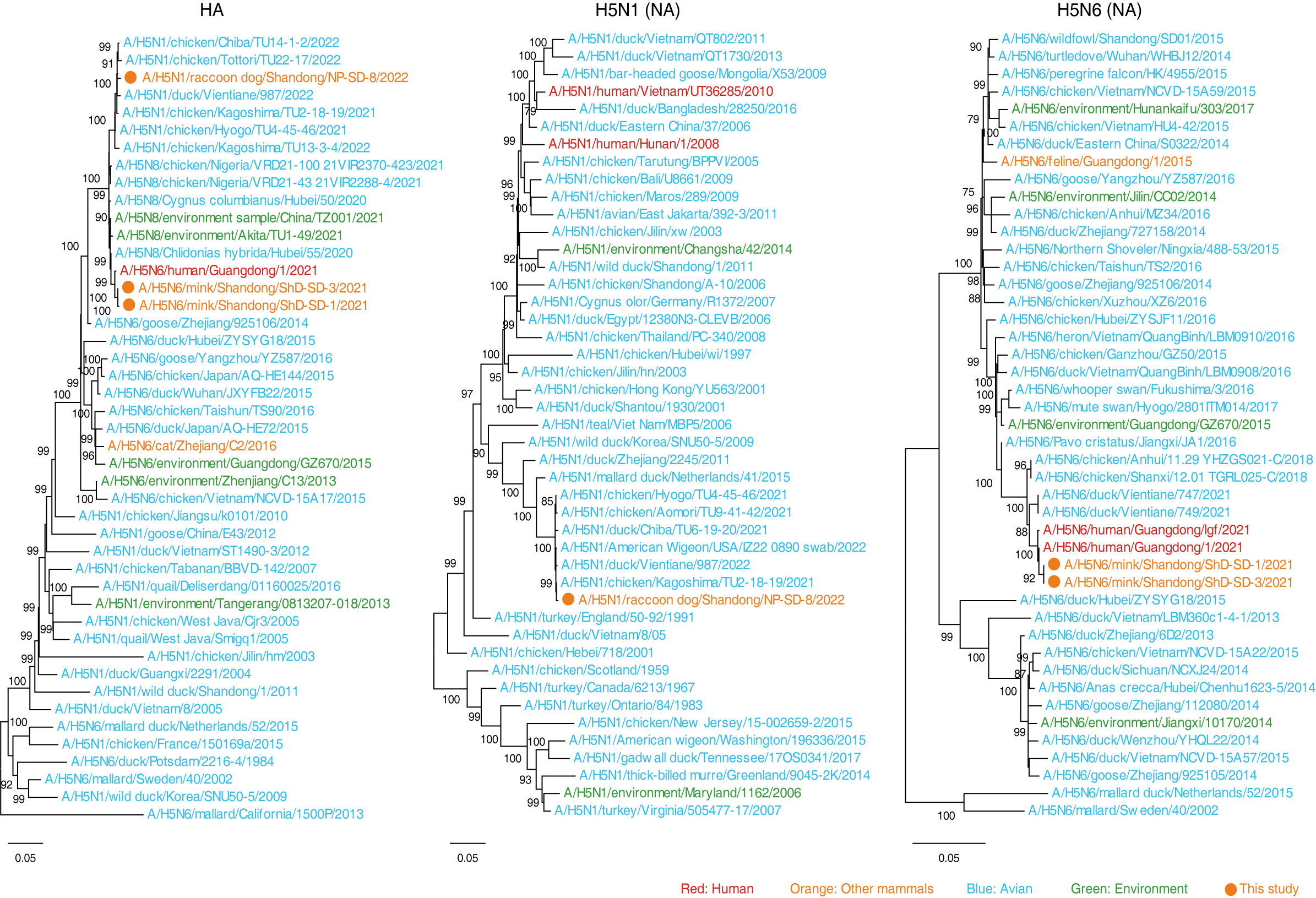

### Extended Data Fig. 11

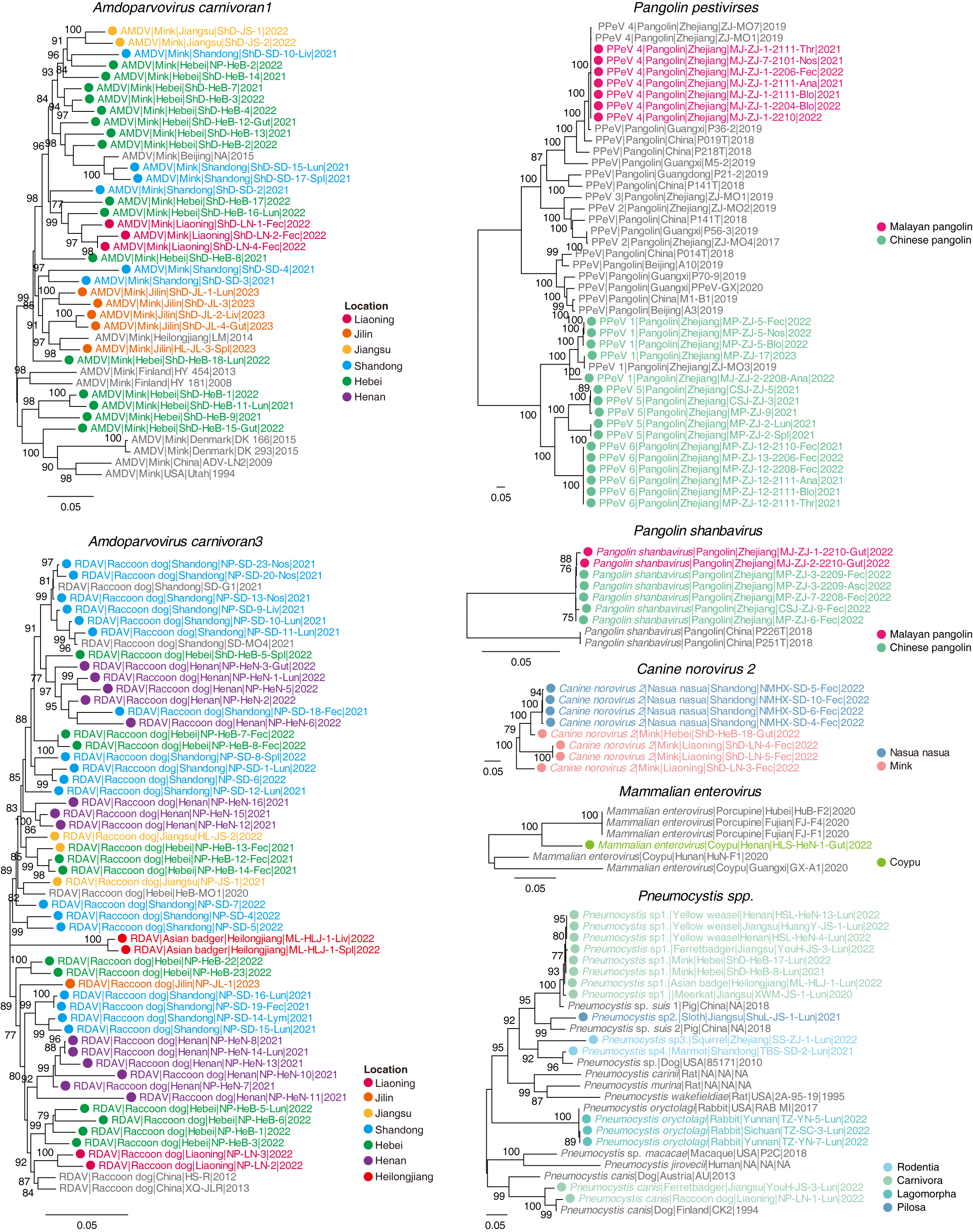
